# Development of an automated high-content immuno-fluorescence assay of pSmads quantification: proof-of-concept with drugs inhibiting the BMP/TGFβ pathways

**DOI:** 10.1101/2023.12.14.571626

**Authors:** Valia Khodr, Laura Clauzier, Paul Machillot, Adrià Sales, Elisa Migliorini, Catherine Picart

**Affiliations:** Université Grenoble Alpes, INSERM, CEA, U1292 Biosanté, CNRS EMR BRM 5000, 17 rue des Martyrs, 38054 Grenoble cedex, France; CNRS, Grenoble Institute of Technology, LMGP, UMR 5628, 3 Parvis Louis Néel, 38016 Grenoble

**Author notes:** co-corresponding authors: Catherine Picart, Valia Khodr. Co-first authors (Valia Khodr, Laura Clauzier, Catherine Picart).

**Keywords:** High-content screening, Immunofluorescence, BMP, BMPR, drugs, biomaterials

## Abstract

Bone morphogenetic proteins (BMPs) and transforming growth factors (TGF-β) are members of the transforming growth factors superfamily, known for their role in several physiological and pathological processes. These factors are known to bind *in vivo* to BMP and TGF-β receptors respectively, which induces the phosphorylation of the Smad (pSmad) transcription factors. This pathway is generally studied with western blot and luciferase bioluminescence assay, which present some limitations. In our work, we developed and optimized a high-content immunofluorescence assay to study the pSmad pathway on glass as well as on biomaterials by overcoming the technical challenges raised by image acquisition and analysis. Furthermore, with this assay, we present here a proof-of concept for drug testing on glass and on biomimetic films using drug inhibitors of the BMP receptor and of the TGF-β receptors. Altogether, our results open perspectives for future drug testing on biomimetic films that present various growth factors and extracellular matrix proteins or peptides.

## Introduction

The transforming growth (TGF-β) superfamily is a large group of structurally related growth factors (GF) grouped into different families, notably the bone morphogenetic (BMP) and TGF-β families. These GF have been reported to have various physiological roles in development and organ regeneration ((1). They have been described to signal *in vivo*, by binding to a ternary complex of type-I and type-II BMP or TGF-β receptor (called BMPR and TGF-βR respectively), which then activates a cascade of signaling pathways such as the canonical Smad pathway and other Smad-independent pathways. In the Smad pathway, the ternary complex of GF and receptors activates Smad 1/5/9 in response to BMP-2, and the Smad 2/3 with TGF-β, by undergoing a carboxy-terminal phosphorylation from the type-I receptor kinase (2). The phosphorylated Smads translocate and accumulate in the nucleus (3) and induce the transcription of several regulation genes (4).

The fact that the pSmad signaling is activated by BMP that has a major osteogenic role *in vivo* and *in vitro*, led to the consideration of Smad as an essential marker of early bone regeneration (5). Consequently, this pathway was extensively studied by the standard molecular techniques such as western blot, gene reporter assay under a specific promoter (like bioluminescence luciferase), and immunofluorescence. The western blot technique has the advantage of having a high sensitivity of the picogram order, and being selective in binding specific target antibody in the presence of heterogeneous mixture of several proteins (6). Similarly the gene reporter based on bioluminescence luciferase technique is characterized with a high sensitivity of the order of 10 femtogram (7). However, these techniques have limitations in terms of their need for optimization steps, variation of signal quantification, in addition to the limitation of the number of samples in the case of western blot. Most importantly, these techniques measure the signal average across a population of cells (6,8). On the other hand, the immunofluorescence technique is generally used for visualization purposes, since the quantification in experiments with several conditions is usually a tedious and lengthy task. However, this technique has the advantage of being compartment-specific and quantitative at a single-cell level. This characteristic could be particularly interesting in detecting phenotypic changes that affect subpopulation of cells (9,10).

The BMP and TGF-β Smad signaling is studied extensively, due to its association with several diseases such as Fibrodysplasia ossificans progressive, Barret esophagus, leukemia and fibrosis (11). Thus, pharmaceutical companies have been focusing on the discovery of new target molecules that can modulate their signaling, in an effort to develop new therapies. The modulation of BMP and TGF-β signaling can be done at different steps of their signaling, but it is mainly the receptor kinase inhibitors that have been developed (12). Thus, our aim was first to develop a high-content screening assay to quantify pSmad intensity on the standard glass condition. Our second aim was to apply this assay to thin biomimetic coatings made of polylysine/hyaluronic acid (PLL/HA) previously developed in the team (13,14), and to optimize it to arise above the difficulties of doing cellular studies on biomaterials (15). Indeed, performing drug screening on biomaterials is not common, yet it has been recently applied to develop a human breast cancer model and hence investigate the effect of extracellular matrix in inducing drug resistance (16). In that context, drug tests on biomaterials could reveal notable processes that could be masked on glass surfaces. Here, using C2C12 skeletal myoblast as a reference model for BMP-responsive cells (17), we show a proof-of-concept with the receptor-kinase inhibitor drugs LDN-193189, inhibitor of the BMP type 1 receptors (ALK1, ALK2, ALK3, ALK6), as well as Galunisertib and Vactosertib, inhibitors of the TGFß type I receptor ALK5, on glass and biomaterials using the fluorescence-based pSmad assay.

## Materials & Methods

### Materials and reagents

Poly(ethylene imine) (PEI) and Poly(L-lysine) hydrobromide (PLL) were purchased from Sigma-Alrich (St Quentin Fallavier, France), and Sodium Hyaluronate (HA) from Lifecore medical (USA). Paraformaldehyde (PFA), methanol and acetone were purchased from Sigma Aldrich (Missouri, USA). The rinsing solutions 0.15 M NaCl pH 6.4 and pH 5.5, as well as 1 mM HCL pH 3.7 and 0,15 M NaCl 20 mM HEPES pH 7.4 were all prepared in house. BMP-2 were bought from Bioventus (France) and TGF-β1 from peprotech (France). LDN-193189, Galunisertib and Vactosertib were obtained from Selleckchem (USA). The glass (ref 655891), TCPS (ref 655986), 96-well cell culture microplates, and CellView (10-well cell culture plates, ref: 543979) were purchased from Greiner bio-one (Germany). The 4′,6-diamidino-2-phenylindole (DAPI) was acquired from Invitrogen, Thermofisher Scientific, France. Trihydrate trichlorhydrate de ‘-[4-éthoxyphényl]-5-[4-méthyl-1-pipérazinyl]-2,5 ‘-bi-1H-benzimidazole (Hoechst 33342), anti-pSmad 1/5/9 (ref: 13820S) and anti-pSmad 2 (ref: 18338S) antibodies were obtained from Cell signaling. The secondary antibody conjugated to Alexa555 or Alexa488 were purchased from Invitrogen, Thermofisher Scientific (France).

### Film preparation

The films were prepared as previously described using a modified robot-liquid handling robot (TECAN Freedom EVO® 100, Tecan France, Lyon) in 96-well cell culture microplates with a layer-by-layer deposition of PLL and HA polyelectrolytes (13,14). They were then crosslinked with 1-Ethyl-3-(3-Dimethylamino-propyl)Carbodiimide (EDC) at a concentration of 70 mg/mL and N-Hydrosulfosuccinimide sodium salt (Sulfo-NHS) at a concentration of 11 mg/mL. The films were rinsed with a solution of HEPES NaCl at pH 7, as previously described (18). The bound BMP-2, hereby called bBMP-2, and bound TGF-β, called bTGF-β1, were then loaded in a solution of HCl at a concentration of 20 µg/mL and 0.75 µg/mL respectively, for 2 h at 37°C, followed by six washes with an HEPES NaCl pH 7 solution. Before usage, the films were sterilized by UV for 30 min.

### Cell culture

C2C12 skeletal myoblasts cells, obtained from the American Type Culture Collection, ATCC, were cultured in 1:1 Dulbecco ‘s Modified Eagle Medium (DMEM):Ham ‘s F12 medium (Gibco, Life Technologies, France) supplemented with 10% fetal bovine serum in an incubator at 37°C and 5% CO2. The cells were cultured up to passage 12. They were then seeded at a density of 10 000 cells/cm^2^ in each well.

### Dose response and kinetic studies

The dose response assays consisted of using a range of concentrations of 0.001 pg/mL to 100 ng/mL for sBMP-2, and 0.001 to 1000 ng/mL for sTGF-β1. We used a concentration range of 0.001 pg/mL to 1000 ng/mL of BMP-2 and TGF-β1 on film and glass, with a volume of 50 µL and 200 µL per well, respectively. In both cases, the cells were exposed to the ligands for 1h. After quantification of pSmad values, the EC50 values were obtained by performing a non-linear dose-response fit on Origin. The kinetic studies consisted on using 400 ng/mL of BMP-2 and TGF-β1 for the soluble conditions, and 20 µg/mL for BMP-2 and 0.75 µg/mL for TGF-β1 in the bound conditions.

### Kinetics and drug assay

These assays consisted on seeding the cells with either sBMP-2 or sTGF-β1 at a concentration of 400 ng/mL and 10 ng/mL respectively, on glass microplates, or with the bBMP-2 and bTGF-β1 at a concentration of 20 µg/mL and 0.75 µg/mL respectively, on the biomimetic films. In the kinetic assay, the cells were fixed at different time intervals ranging from 15 min until 3 days. We first exposed the cells to the BMP/TGF ligand for 15 min. Then the drug was added for 30 min, at a concentration range of 0.1 pM to 0.1 µM for LDN-193189 and 10 pM to 10 µM M for galunisertib and vactosetib. The IC50 values were obtained by performing a non-linear dose-response fit using Origin software with the equation 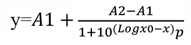, with A1 being the highest value, A2 the lowest, and *p* the hill slope.

### Immunofluorescence assay

The cells were fixed with 4% PFA, and stained with 4′,6-diamidino-2-phenylindole (DAPI) or Hoechst 33342 for the nuclear staining. The pSmad was stained by using a 1/800 dilution of anti-pSmad antibody, and a secondary antibody conjugated to either Alexa Fluor 488 or Alexa Fluor 555 diluted at 1:500 in BSA 3% w/v in PBS.

### Image acquisition

The images were acquired using In Cell GE INCA 2500 imaging system (General Electrics Healthcare, France) with a 20X objective. Similar exposure times were applied for both pSmad1/5/9 and pSmad2. Sixteen to 21 images distributed over each well were acquired and at least 500 cells were analyzed per well. For image analysis, the InCarta® software (General Electrics Healthcare, USA) was used to automatically segment nuclei and cells, and to quantify the nuclear pSmad signal intensity.

### Data analysis

The raw intensity data were normalized by the maximum signal measured, and transformed into a percentage of Smad signaling where the background signal corresponds to 0% response. Reported pSmad intensity values are obtained by averaging the values obtained from at least two biological replicates, and three technical replicates for each experiment. The reported errors are presented as the standard deviation.

## Results and discussion

### Immunofluorescence pSmad assay and image analysis

In this study, we developed and optimized a high-content screening immunofluorescence assay to quantify the translocation of Smad to the nucleus, first on glass as cell culture substrate, then on the biomimetic film coating. We started by optimizing the experimental conditions, notably cell culture, image acquisition and subsequent analysis (**FIG. 1**) to be compatible with both our biomimetic films and the standard glass condition used for imaging (19). For simplicity, sBMP-2 and sTGF-β1 designations will be used to indicate the soluble proteins used on glass surfaces, while bBMP-2 and bTGF-β1 will be used to designate the matrix-bound proteins loaded in the biomimetic films. For image acquisition and analysis, we used an automated imaging system (In Cell analyzer) adapted for high-content screening, as described in the materials and methods section. To quantify pSmad 1/5/9 in the nucleus, the raw image of the nuclei stained with DAPI was used to set a threshold in order to remove the background fluorescence and to select solely the nuclei. This analysis generates a “mask image” of the nuclei, which can be subtracted afterwards from the image of pSmad staining. As a result, a new image is created that shows solely pSmad staining inside the nucleus **(FIG.1.A)**. The nuclear intensity was then quantified using either InCarta®, which is the commercialized program of In Cell analyzer, or can be alteratively analyzed using the widely-used image analysis software ImageJ.

**FIG 1.**
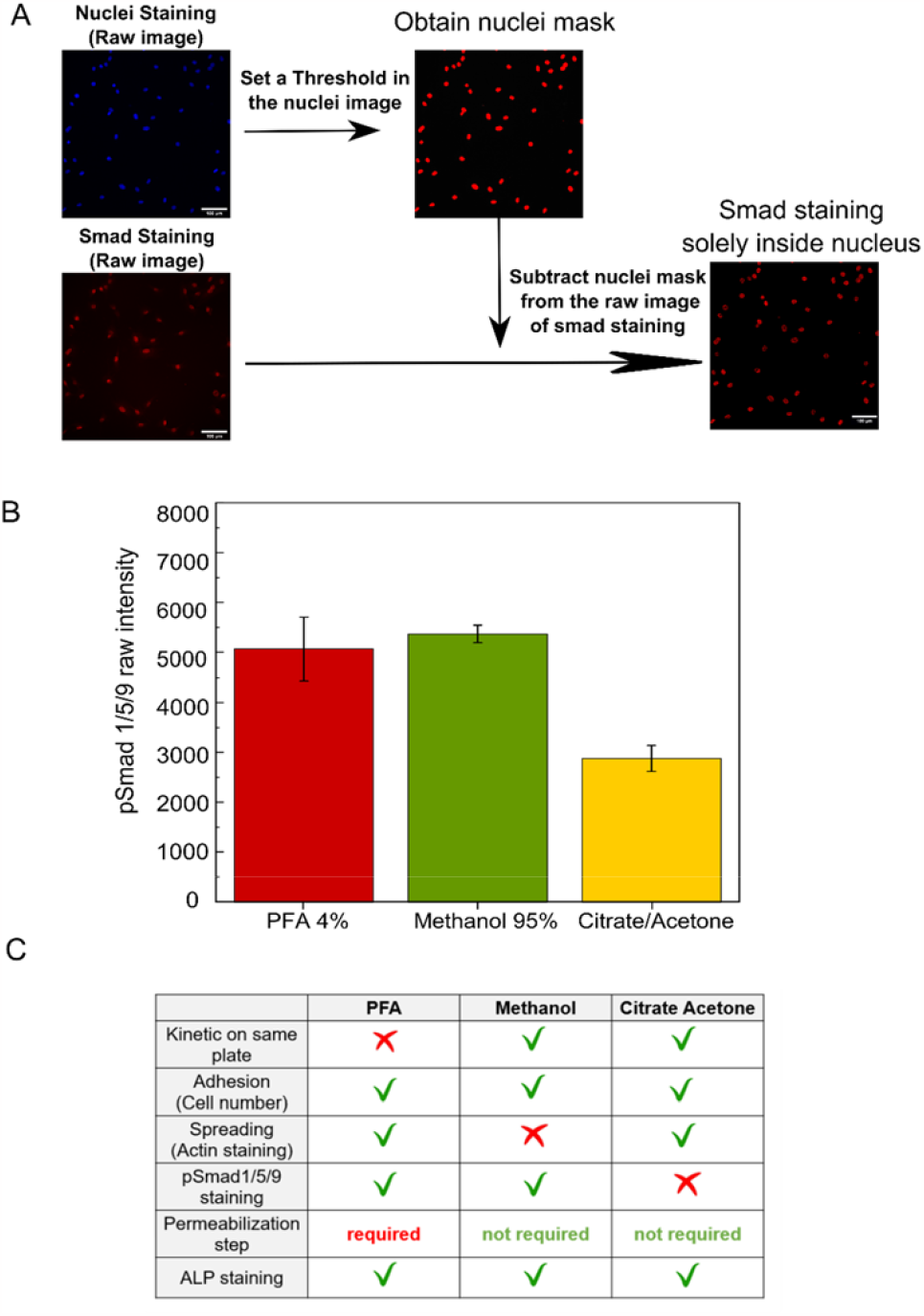
Principle of quantification of pSmad1/5/9 staining in the nucleus and selection of the fixation method for optimized pSmad quantification of parameters related to cell differentiation (pSmad, ALP) and cell adhesion (cell number and spreading area) A). The acquisition was done with an automated inCell analyzer system. The image analysis consisted on first setting a threshold on the raw image of the nuclei, which then creates a mask image that could be subtracted from the raw image of pSmad staining. The resulting image presents then the Smad staining in the nucleus. Scale bar =100 µm. B) Quantification of pSmad 1/5/9 signal in response to sBMP-2, after cell fixation with either 4% PFA, 95% Methanol and 40 mM Citrate/60% Acetone. C) A table summarizing the compatibility of the different fixation solutions with the output parameters studied: kinetics on same plate, adhesion, pSMAD1/5/9 staining, need for a permealization step and ALP staining.

In a kinetic imuno-fluorescence assay, the cell fixation step is essential since cells need to be fixed at different timpe points throughout the experiment. Hence, we first optimized the experimental conditions using soluble BMP-2 (sBMP-2) and compared three different methods of cell fixation: 4% Paraformaldehyde (PFA) that is a commonly-used fixation protocol, 95% methanol and 40 mM citrate/ 60% acetone **(FIG.1.B)**. Immunofluorescence of pSmad 1/5/9 showed that pSmad signal was the highest for fixation with PFA4% and 95% methanol, in comparison to the citrate/acetone mixture. Next, we looked at several other parameters related to i) the possibility to do a kinetic experiment on the same plate, ii) to quantify cell adhesion via the cell number (nucleus labeling) and cell spreading area (actin staining of the cell cytoskeleton), iii) the need for a permeabilization step, iv) the the possibility to perform an alkaline phosphatase (ALP) staining, since it is considered an early marker of differentiation in bone cells (17). The results are summarized in the table of **FIG.1.C**. Overall, a toxicity was found when using PFA for the kinetic assay on the same plate. We observed that the citrate/acetate solution is not compatible with pSmad staining, as also seen with the lower Smad intensity level **(FIG.1.A)**. In contrary, fixation with 95% methanol was compatible with pSmad staining and did not require a permeabilization step, but it affected the actin staining. Thus, we conclude that both PFA and methanol can be used to study pSmad signaling depending on the desired readout, PFA fixation being well adapted for studies at a single time-point, while methanol can be used for kinetic studies, but it is not suited for quantitative measurement of cell spreading area.

After defining the fixation conditions, we needed to optimize the imaging conditions, especially the type of antibodies for cells grown on the different types of supports (glass or tissue culture polystyrene, TCPS) without or with a film coating (**FIG. 2**). We first compared the effect of two different fluorescent primary antibodies Alexa488 and Alexa555, and of two supports (glass, TCPS) on pSmad fluorescence intensity **(FIG.2.A-B)**. pSmad 1/5/9 signaling was analyed for cells cultured for 1h with sBMP-2 at 400 ng/mL. We observed a higher pSmad intensity **(FIG.2.A)** and a lower background intensity **(FIG.2.B)** with Alexa555 in comparison to Alexa488. Moreover, the background intensity was lower on glass plates, which were chosen for their image clarity. Notably, the background intensity on films was lower when Alexa555 antibody was used for pSmad staining. As last step, we investigated the effect of the dilution level of the antibodies used for immunofluorescence. We thus used five dilutions of pSmad 1/5/9 primary antibody from 1/400 to 1/1200 and two dilutions of Alexa 555 secondary antibody ( 1/500 and 1/700) **(FIG.2.C)**. While we observed negligible differences in secondary antibody, the concentration of primary antibody slightly impacted the signal levels. Therefore, we selected an intermediate dilusion of 1/800 of primary antibody and a 1/500 dilution of secondary antibody as optimal dilutions, to perform the immunofluorescence test.

**FIG 2.**
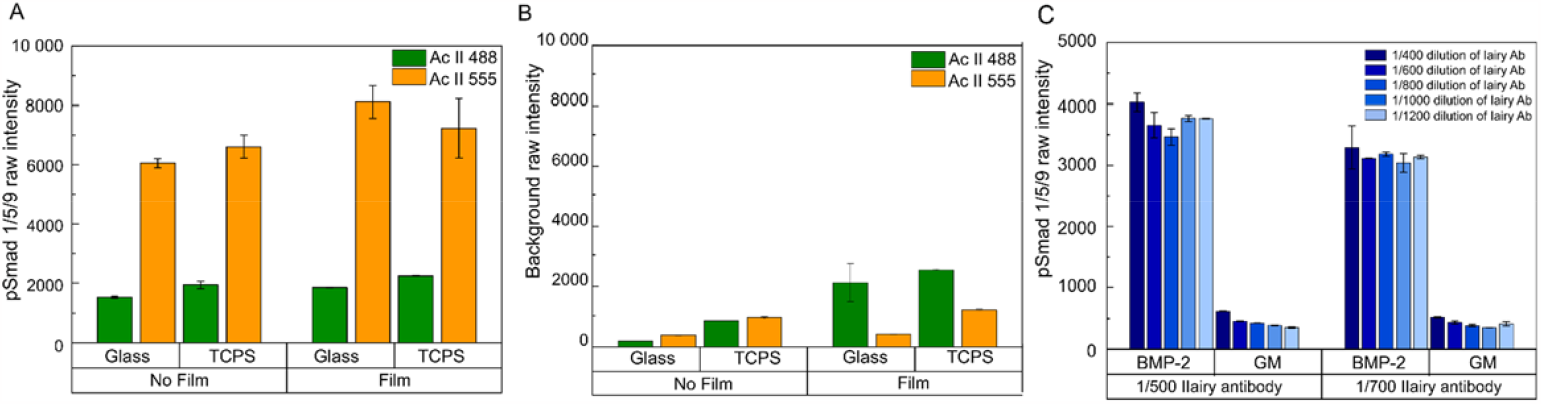
Optimization of antibodies on the different types of plates. The plates were either made of glass, TCPS, and were without or with a biomimetic film coating. A) The pSmad 1/5/9 intensity was measured with the secondary antibody Alexa488 and Alexa555 without and with film, on glass and TCPS. B) The background intensity was measured in the same experimental conditions. C) The pSmad 1/5/9 intensity was measured with 1/400, 1/600, 1/800, 1/1000 and 1/1200 dilutions of pSmad 1/5/9 primary antibody, as well as with 1/500 and 1/700 dilutions with the secondary Alexa555 antibody, in the presence and absence of BMP-2.

After selecting the preparation and imaging conditions, we performed a dose-response assay by adding sBMP-2 and sTGF-β1 to C2C12 cells that were pre-cultured for 24h (**FIG 3**). After 1h of contact with sBMP-2 or sTGF-β1, the cells were fixed (**FIG 3.A.)**. Cells in contact with sBMP-2 were stained for pSmad 1/5/9 while those in contact with sTGF-β1 were stained for pSmad2. As seen in **FIG.3B**, a clear increase in intensity of pSmad 1/5/9 in the nucleus was visible when the concentration of sBMP-2 increased from 10^-1^ to 10^4^ ng/mL. The control condition without BMP-2 (GM) was effectively negative. The pSmad intensity in the nucleus was subsequently quantified and the half-maximal effective concentration (EC50) calculated, after performing a non-linear dose-response fit **(FIG.3.C)**. The EC50 was determined to be 0.92 ± 0.06 ng/mL for sBMP-2(equivalent to 0.2 ± 0.01 ng, when expressed in absolute mass) and 0.12 ± 0.01 ng/mL (corresponding to 0.02± 0.002 ng) for sTGF-β1.

**FIG 3.**
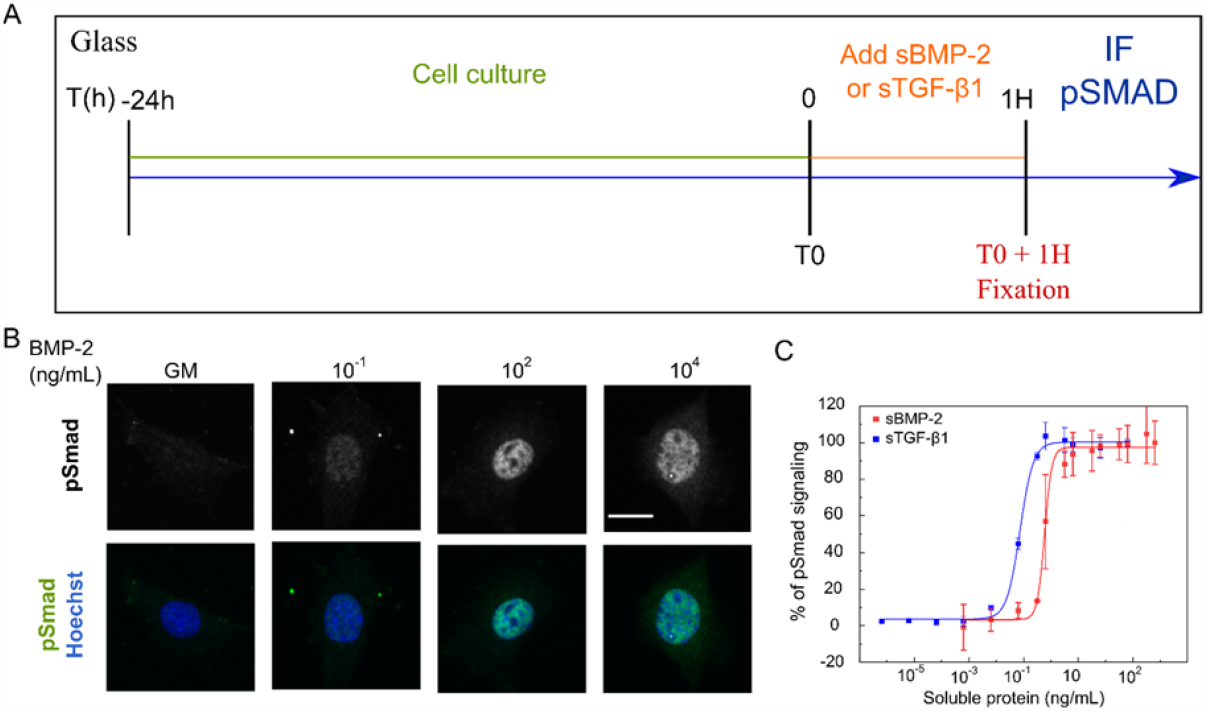
A dose-response immunofluorescence assay for pSmad nuclear translocation. A dose response assay was performed. A) The experiment consisted on seeding the cells for 24h and then adding sBMP-2 and TGF-β1 at a concentration of 10^-6^ to 10^3^ ng/mL. After 1h of protein stimulation, the C2C12 cells were fixed and immunostained using Hoechst and pSmad 1/5/9 or pSmad 2 antibody. B) Cellular images of the dose-response assay with BMP-2, with pSmad 1/5/9 and nuclei staining. Scale bar= 26 µm. C) The dose response curves with soluble BMP-2 (sBMP-2) and TGF-β1 (sTGF-β1).

These values are consistent with the literature data since a study reported the EC50 values, calculated using an AlphaLisa test in C2C12 to be 0.9 ± 0.1 ng/mL for BMP-2 and 0.052 ± 0.002 ng/mL for TGF-β1 (20). In addition, other studies determined using luciferase assay, the EC50 of BMP-4 to be 0.3 ng/mL in human cervical carcinoma cells (21), and 2.4 ng/mL in human embryonic kidney cells (HEK293) (22). If compared to BMP-4, the values of EC50 for BMP-2 appears to be comparable to literature data.

### Kinetic of pSmad phosphorylation with soluble and matrix-bound BMP-2 and TGF-β1

After defining the optimal experimental parameters, we examined the kinetics of pSmad signaling with BMP-2 and TGF-β1 on both glass and film with the aim of defining the optimal time-window to perform the drug experiments. The kinetic experiments were done for cells cultured on glass plate with sBMP-2 and sTGF-β1, and for cells cultured on the biomimetic films with bBMP-2 and bTGF-β1. The cells were fixed at different time points ranging from 30 min up to 3 days and pSmads were quantified at each time points (**FIG.4.A-B)**.

The pSmad intensities exhibited differences depending on the type of ligand (BMP-2 versus TGF-β1) and presentation mode of the ligand (soluble versus matrix-bound). For sBMP-2, pSmad1/5/9 intensity was high at 30 min, increased and reached a peak intensity at 1.5 h before decreasing (**Fig 4.A**). A second peak could be observed at 3h. Then pSmad nuclear signal remained sustained at 40% up to 24h, before retaining to basal level at 48h.

**FIG 4.**
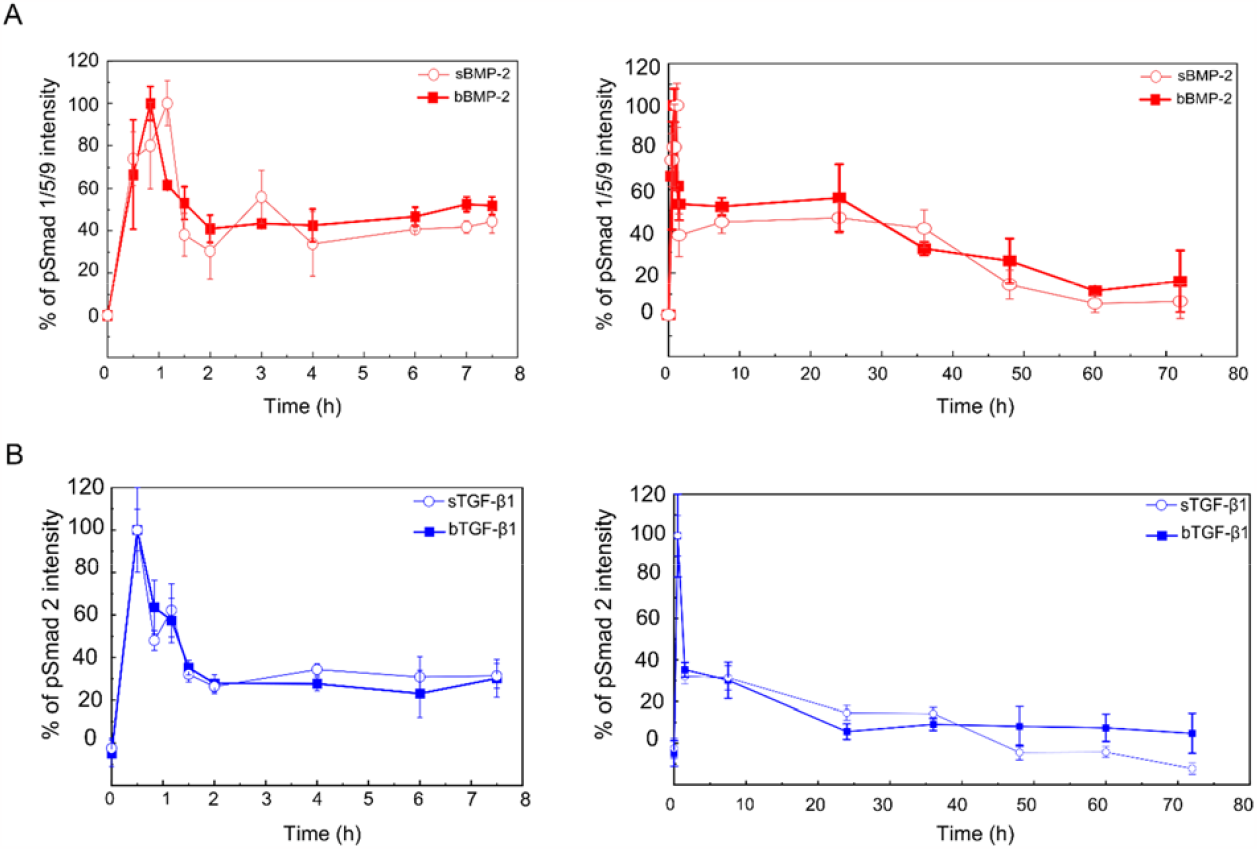
Kinetic study of pSmad phosphorylation in C12C12 skeletal myoblasts in the presence of soluble and matrix-bound BMP-2 and TGF-β1. Data points corresponding to the cell experiments on glass are represented with an empty circle, while those on the biomimetic films are represented with a filled rectangle.A) A kinetic graph of pSmad 1/5/9 after 1h of C2C12 cell exposure to BMP-2 on glass and film. B) A kinetic graph of pSmad 2 after 1h of C2C12 cell exposure to TGF-β1 on glass and film.

For cells on bBMP-2, the pSmad peak was sharp at ∼1h, before decreasing to ∼50% and remaining sustained up to 24h. No second pSmad peak was observed. After 24h, it gradually decreased until day 3 where the signal was still close to 20%.

In contrast, for TGF-β **(FIG.2.B)**, the pSmad2 kinetics was faster with an initial peak at 30 min for sTGF-β and a second peak at 1h15 before quickly returning to a low level at ∼25% at 1h30. Then, the signal fully returned to its basal level at 48h, some minor oscillations being visible over time.For bTGF-β, the first peak was at the same time point a for sTGF-β, but the second peak was less marked. A decrease to 25% was also fast and quickly observed at 1h30, then the signal was sustained until 8h, after which it returned to its basal level.

These results are compatible with the literature as kinetic studies of BMP-2 done by western blot showed the presence of a pSmad 1/5/8 signal after 30 min with a sustained signal up to 180 min on different types of biomaterials (23–25). Similarly, studies with TGF-β1 in different cell models showed the activation of pSmad 2 signaling after 30 min of TGF-β1 addition, and up to 8h in continuous ligand stimulation (26,27).

The presence of a sustained signal may be explained by the mechanism of Smad regulation of import and export to/from the nucleus. Indeed, it was reported that Smad 3 possess an NLS-like sequence in the MH1 domain, capable of binding a karyopherin (the importing β1) and hence induce its import to the nucleus (28). In contrast, Smad 2 seemed to be by directly interacting with the nucleoporins to translocate to the nucleus (29).

To conclude, kinetics of pSmad1/5/9 and pSmad 2 exibit some distinct differences, while the presence of the films contributes to dampen the second peak that is clearly observed for soluble ligands, and to sustain signalling.

Since we observed an initial peak varying from 30 min to 1h30 for both BMP-2 and TGF-β1 proteins for cells either cultured on glass or on the films, we selected 1h as common time point for the future experiments using drugs.

### Test of drug inhibitors against BMP receptors for cells cultured on glass and on biomaterials

After optimizing the time frame, we performed dose-response assays with soluble and matrix-bound proteins to determine the optimal concentration for the drug experiments.

As a first control, since the drugs are in DMSO, we tested the effect of DMSO on the fluorescence intensity of Smad 1/5/9 and Smad 2/3 **(SI.FIG.1-A-B)**. We did not observe any effect on the Smad intensity signal for levels of DMSO up to 1%, which was the maximal concentration used in our assay, while 2% and 5% levels showed an important increase in Smad signaling.

For the film condition, we seeded the C2C12 cells directly on the biomimetic films loaded with bBMP-2 and bTGF-β1, and on the glass condition supplemented with sBMP-2 or sTGF-β1 simultaneously as described in **FIG.3.A**. After quantification of Smad intensity **(FIG.3.B-C)**, we determined the EC50 values, which were similar for the glass and film conditions. The values of EC50 on glass are 120 ± 47 ng/mL (corresponding to 24.0 ± 9.4 ng) for sBMP-2, and 0.5 ± 0.16 ng/mL (equivalent to 0.1 ± 0.03 ng) for sTGF-β1. As for the film, the EC50 values were determined to be 288 ± 20 ng/mL (equivalent to 14.4 ± 1.0 ng) for bBMP-2, and 8 ± 2 ng/mL (corresponding to 0.4 ± 0.1 ng) for bTGF-β1. We note that the EC50 value of sBMP-2 when cell seeding and the protein is added simultaneously **(FIG.5.B-C)** are higher than the EC50 value of sBMP-2 and sTGF-β1 when cell have adhered to 24h (0.2 ± 0.01 ng and 0.02 ± 0.002 ng, respectively) **(FIG.3.A)**.

**FIG 5.**
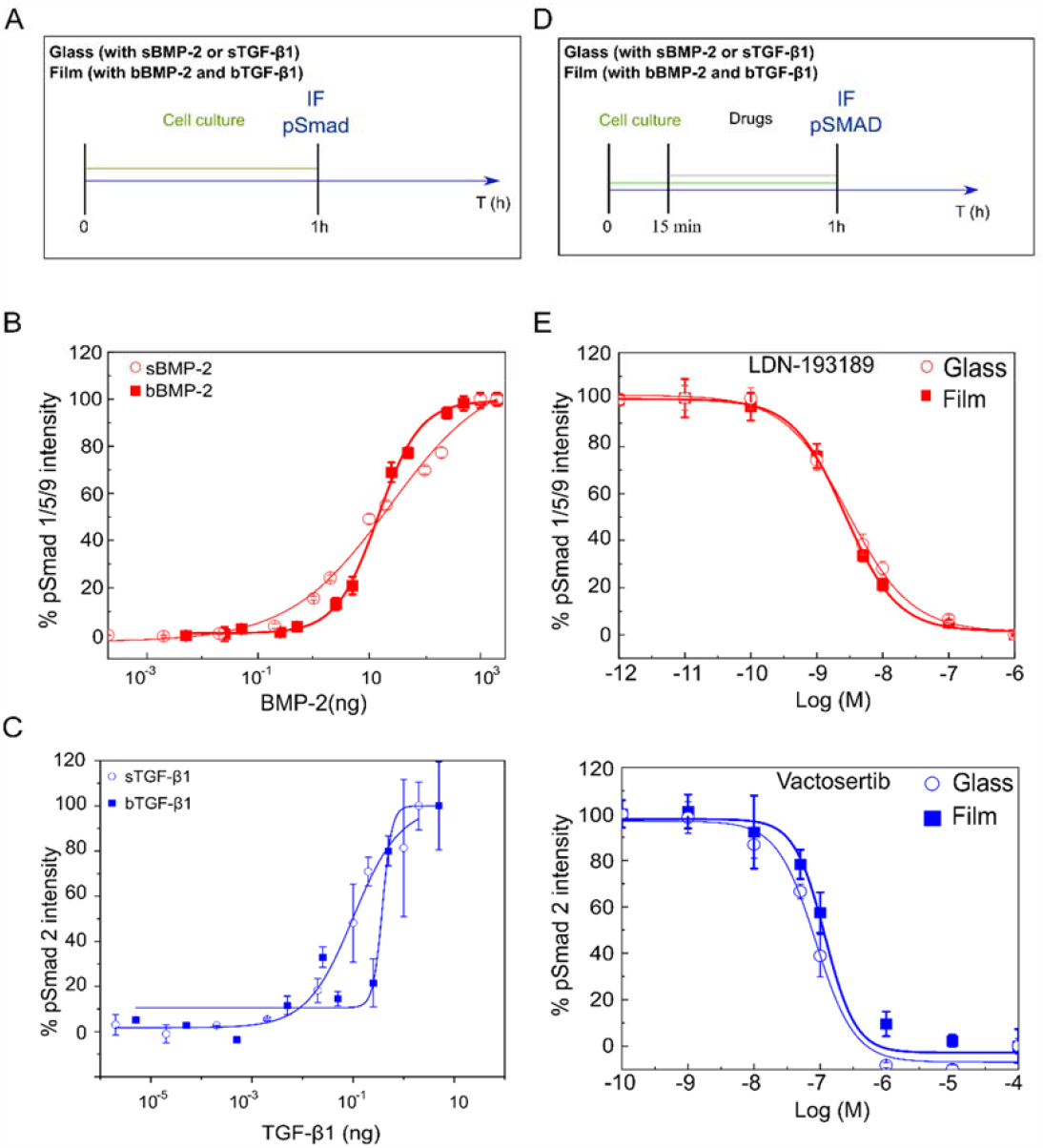
ALK inhibitor drug assay with soluble and matrix-bound BMP-2 and TGF-β1. A) Schematics of the experimental protocol of the dose-response tests on glass and biomaterials with sBMP-2/bBMP-2 and sTGF-β1/bTGB-β1. B) A dose-response test with bBMP-2 and sBMP-2 on film and glass respectively with C2C12 cells. C) A dose-response test with bTGF-β1 and sTGF-β1 on film and glass respectively with C2C12 cells. D) Schematics of the experimental protocol of the drug tests on glass and biomaterials with sBMP-2/bBMP-2 and sTGF-β1/bTGF-β1. E) Drug assay of LDN-193189 with sBMP-2 and bBMP-2, and F) Vactosertib with sTGF-β1 and bTGF-β1 with C2C12 cells on glass and films.

These IC 50 values are in agreement with previous studies where the adhesion of C2C12 on the biomimetic films with bBMP-2 was studied (30). They showed that the presentation of the usage of biomimetic films and presenting BMP in a matrix-bound manner leads to a crosstalk between the BMPR and integrins that are responsible for the adhesion. They also showed that the cascade of events starts with BMP-2 binding to BMPR followed by integrin activation, before the Smad signaling (30). We can thus hypothesize that the added BMP-2 when the cells have not yet adhered, are used in focal adhesion formation to ensure adhesion, in addition to the activation of BMPR to initiate pSmad signaling. We can also assume that a similar process involving integrin occurs when cells are seeded on glass. Hence, the adhesion process could explain the need for a higher quantity of protein when cells have not yet adhered in comparison to when the cells have already adhered.

After characterizing the cellular response to BMP-2 and TGF-β1 on biomaterials, we chose suitable concentrations higher than the EC50 values to do the drug experiments. We selected LDN-193189, one of the first discovered kinase inhibitor of Type I BMP receptors (31), in addition to two ALK5 kinase inhibitors currently being developed in clinical trials for cancer: Galunisertib, which was stopped at phase 2 of clinical trials by the sponsor company, and Vactosertib which is currently in phase 2 (32,33). The experiments were performed as shown in **FIG.5.D**. The drugs were added for 45 min, followed by cell fixation.

After quantification of the Smad signal, we were able to determine the half-maximal inhibitory concentration (IC50). The results showed similar curves and IC50 values for both the film and glass conditions **(FIG.5.E)**. The IC50 for LDN-193189 is determined at 3.0 ± 0.03 nM and 2.7 ± 0.2 nM for glass and film respectively. While the IC50 of Vactosertib are 85 ± 12 nM and 118 ± 21 nM and Galunisertib are 1276 ± 79 nM and 1587 ± 552 nM **(SI.FIG.2)**, and for glass and film respectively.

In order to validate that our IC50 are not BMP concentration-dependent, we performed a drug assay with LDN-193189 and several concentrations of BMP-2. The results showed that the pSmad intensity is independent of the BMP-2 concentration **(SI.FIG.3)**. This result may be explained by the action mode of the inhibitors that inhibits the BMP and TGF-β signaling at the receptor level.

The data in the literature for the exact IC50 varies, as it is cellular and assay dependent. For instance, the manufacturer ‘s reported IC50 values are 0.8 nM, 11 nM, and 56 nM for LDN-193189, Vactosertib, Galunisertib and respectively using *in vitro* kinase assays, commonly used in drug development. This kinase assay consists in measuring the kinase activity by determining the amount of radioactive ATP transferred to the substrate (34–37). Furthermore, a study of pSmad 1/5/9 signaling inhibition with BMP-4 and LDN-193189 determined its IC50 to be 4.9 nM (31). On the other hand, other studies have reported the IC50 of Galunisertib at 50 to 430 nM depending on the used assay (38). Nevertheless, in our work the values of IC50 are similar for both film and glass conditions with the three drugs, indicating the robustness of the immunofluorescence assay and its application in different experimental conditions for soluble and matrix-bound proteins, on glass or on biomaterials. This also confirms that the drug is acting similarly and independently of the presentation mode of BMPs, which indicates that biomaterials could be used in future drug testing.

In order to determine the stability of our assay in performing high-throughput screens, we carried out a Z’ factor experiment, where LDN-193189 was added at a concentration of 100 nM and BMP-2 at 2 µg/mL and 400 ng/mL for the film and glass respectively. The study demonstrated a Z’ factor of 0.3 for glass and 0.6 for film. Since an acceptable assay has a Z’ factor around and higher than 0.5, we show here that our pSmad assay is adapted for high-content drug assays on both glass and biomaterials.

The similarity in IC50 values between the film and glass indicates that the film is comparable to the glass condition, which is commonly used as reference for *in vitro* drug testing. From our study, we can conclude that this optimized pSmad test is adapted to do high content drug screening on biomimetic materials, similarly to the standard glass condition. These results further show the possibility of future studies on biomaterials to investigate pathologies in a biomechanical context by modifying the biomimetics films’ stiffness.

## Conclusion

In this work, we present a high-content immunofluorescence assay to study the pSmad pathway for cells on glass and biomaterials. We show the optimization of the classic protocol to overcome the limitations of doing certain tests such as the kinetic tests on protein-loaded biomaterials. We also demonstrate a proof-of-concept of pSmad activity-based drugs assay on biomaterials. This assay may thus be further used on the biomimetic films containing other ECM proteins or growth factors.

## Acknowledgments

The authors thank Marie-Odile Fauvarque, Emmanuelle Soleilhac and Catherine Pillet for fruitful discussions and advices, notably regarding the drug assay experiments. We are also grateful to Sabine Bailly for her advices, reading of the mansucript, feedback and suggestions.

## Data availability

The data that support the findings of this study are available from the corresponding author upon request.

## Funding and additional information

The study was supported by the Agence Nationale de la Recherche (ANR CODECIDE, ANR-17-CE13-022), and Institut Universitaire de France (IUF) to C.P. In addition to the European Research Council (ERC) under FP7 programm (ERC Biomim GA259370), the ERC POC Bioactive coatings (GA 692924) and by the Fondation Recherche Medicale (FRM, grant DEQ20170336746) to C.P. V.K. was supported by a PhD. fellowship from Grenoble Institute of Technology.

