## supplemental file for "Development of an automated high-content immuno-fluorescence assay of pSmads quantification: proof-of-concept with drugs inhibiting the BMP/TGFβ pathways"

### Khodr et al, Supplementary data

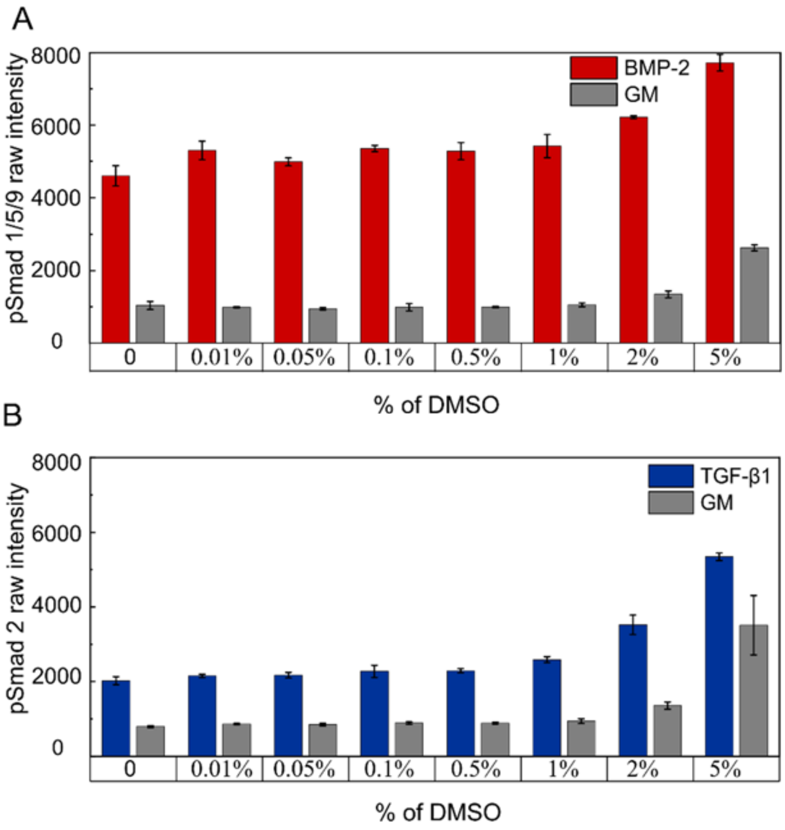


**SI.FIG.1. Control test with DMSO for the drug experiments.** A) The pSmad 1/5/9 raw signal was measured in the presence and absence of BMP-2 with increasing percentage of DMSO up to 5%. B) The pSmad 2 raw signal was measured in the presence and absence of TGF-β1 with increasing percentage of DMSO up to 5%.

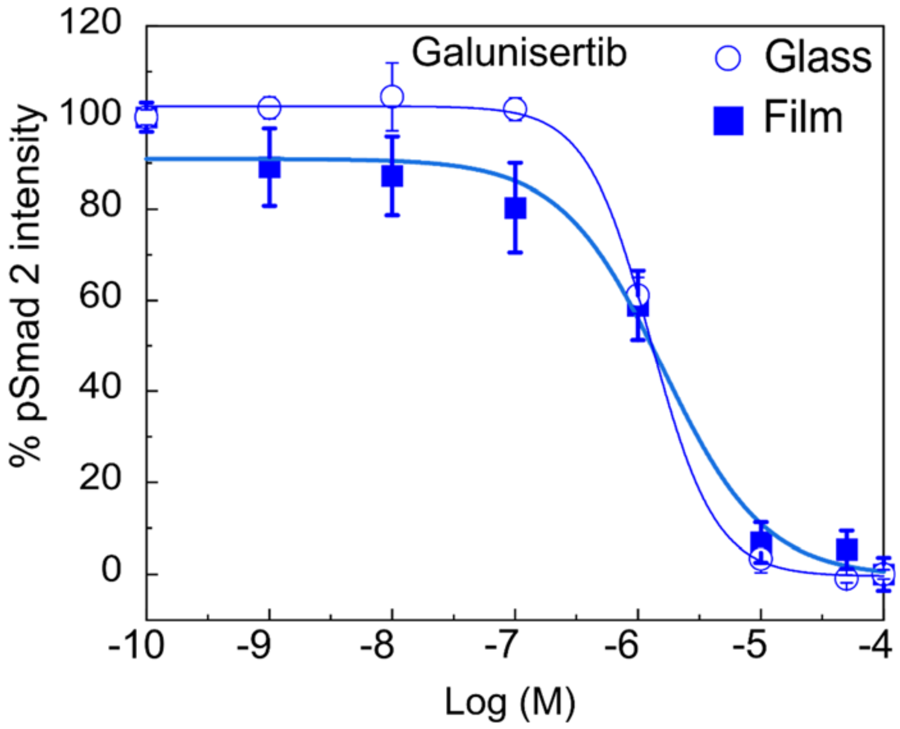


**SI.FIG.2. Drug assay of Galunisertib with TGF-β1 with C2C12 cells on glass and film.**

A dose-response test with bTGFβ1 and sTGFβ1 on film and glass respectively was done with C2C12 cells in the same conditions as in Figure 5.

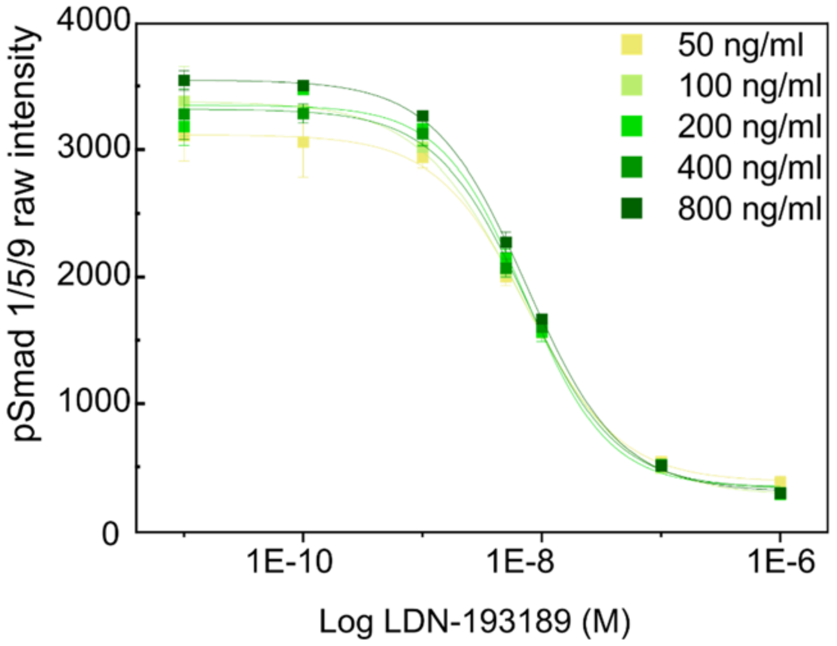


**SI. FIG.3. Test of LDN-193189 in the presence of increasing concentrations of soluble BMP-2.** A drug test was performed with LDN-193189 and different concentrations of BMP-2 in solution (50 ng/mL, 100 ng/mL, 200 ng/mL, 400 ng/mL and 800 ng/mL).
